# Unraveling codon usage of *Escherichia coli* using machine learning

**DOI:** 10.1101/2023.02.07.527422

**Authors:** Bifang Huang, Yunzhuo Hu, Xuanyang Chen, Shiqiang Lin

**Affiliations:** Key Laboratory of Crop Biotechnology, Fujian Agriculture and Forestry University, Fujian Province Universities, Fuzhou, China; College of Life Science, Fujian Agriculture and Forestry University, Fuzhou, China; College of Agronomy, Fujian Agriculture and Forestry University, Fuzhou, China

## Abstract

Machine learning is used to investigate the codon usage of protein-encoding genes, which is one of the fundamental questions of molecular biology. The presentation, parameter learning, and decoding of the conditional random field (CRF) model are implemented and then utilized to analyze the codon usage of the genes of *Escherichia coli* and its phages. Most genes of *E. coli* use codons conforming to the weights of the model determined by all *E. coli* genes. Phages use the codons like their host *E. coli*. Finally, the study evaluates the codon usage of several example genes in the context of the model. These results help to understand the codon usage in *E. coli*.

## Introduction

In 1958, Francis Crick proposed the central dogma of molecular biology, i.e., the genetic information in the cell flows from DNA to RNA and then from RNA to protein (Crick, 1970). The DNA sequence determines the RNA sequence and the RNA sequence determines the protein sequence. However, due to the synonymous codons, there might be a huge number of possible DNA sequences capable of encoding a given protein. This raises the question of how the codons are used for the protein-encoding genes. Reports have shown that codon usage is important for mRNA stability and gene expression level (Zhou et al., 2016, Quax et al., 2015, Kudla et al., 2009). The synonymous mutations are associated with protein folding and human disease (Walsh et al., 2020, Sauna and Kimchi-Sarfaty, 2011). A recent study showed that the synonymous mutations of some genes in yeast are detrimental (Shen et al., 2022). Evidently, the codons are not randomly used in the protein-encoding genes. The problem is how to describe the codon usage, or which model shall be used.

Deep learning seems attractive since it has played important roles in image recognition, protein engineering, and protein structure prediction (Zhong et al., 2021, Alley et al., 2019, Jumper et al., 2021, Baek et al., 2021). Researchers borrowed ideas from natural language processing and developed the protein language model for analyzing large numbers of protein sequences (Elnaggar et al., 2022). However, it is difficult to interpret the learned parameters in a practical sense from the universal architecture of deep learning. Moreover, deep learning needs a large sample to train the magnificently numerous model parameters. Moreover, there are lots of nodes in one layer and many layers within the neuron network of the deep learning model, which needs tens of thousands, or even more parameters for model description. Consequently, a large sample is required to train these parameters. The *E. coli* has only less than 5, 000 genes, by no means an enough large sample for the deep learning model to train the magnificently numerous parameters.

Another approach is the conventional machine learning models that are reasonably mature and interpretable (Greener et al., 2022). The parameters are relatively few and the training does not require a large sample. Among the conventional machine learning models, the conditional random field (CRF) model is excellent for describing the relationship between the protein sequence and the gene sequence as it is essentially the same as the labeling of sequential data (Lafferty et al., 2001). Fig. 1 shows the principle of the conditional field model for describing the codon usage of protein-encoding genes. Calculating the unnormalized probability of each path takes all of the edges and vertices into account, which overcomes the label bias problem and allows long-range dependency. In addition, the cost function of the CRF model during parameter learning is a convex function, which behaves well for allowing an arbitrary approximation to zero in theory. These merits make the CRF model an exceptionally good model for describing codon usage.

**Fig. 1.**
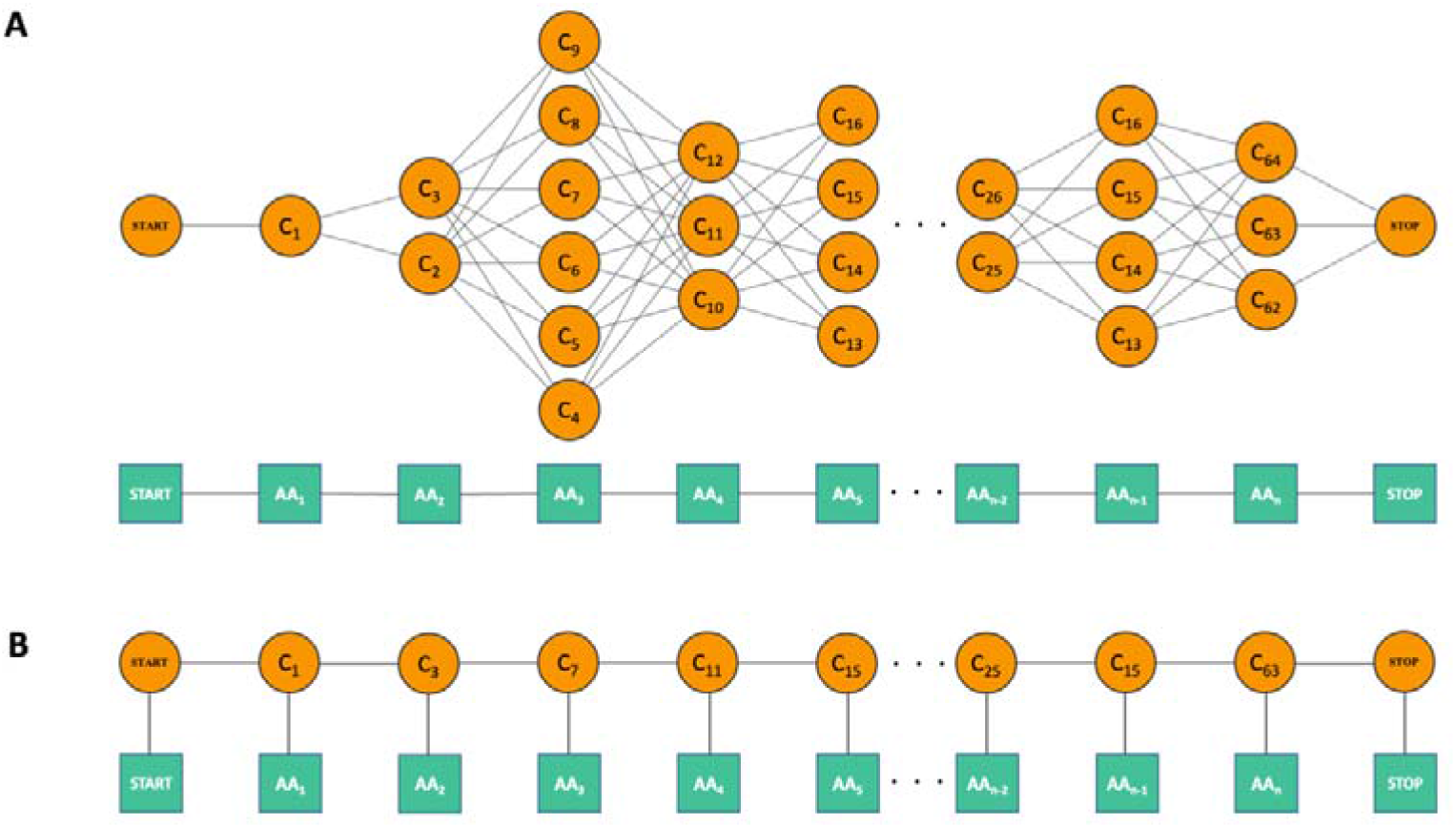
A pictorial representation of the conditional random field model of protein and DNA. (A) All possible DNA sequences for a given protein sequence. (B) One of the possible DNA sequences for a given protein sequence. An amino acid is called an observation and a codon is called a state. A codon-codon pair is called a state pair and a codon-amino acid is called a state observation pair.

For the DNA sequence in (B), the unnormalized probability can be calculated by exp (START-START+START-C_1_+C_1_-AA_1_+C_1_-C_3_+C_3_-AA_2_+,…,+C_15_-C_63_+C_63_-AA_n_+STOP-STOP),where each item inside the parenthesis, C_1_-C_3_ for instance, stands for the weight of C_1_-C_3_, and the unnormalized probability of any other DNA sequence in (A) can be calculated in this way. The normalized probability of each DNA sequence is obtained by dividing its unnormalized probability by the sum of the unnormalized probabilities of all DNAs.

Here, we deal with the genes of *Escherichia coli* str. K-12 substr. MG1655 and write the Python codes for the presentation and parameter learning of the CRF, the analysis of *E. coli* and phage genes, aiming to describe and explain the codon usage of protein-encoding genes, which is of both theoretical and practical importance.

## Results

### Parameter learning and analysis of CRF

The S algorithm is applied to perform the improved iterative scaling (IIS) for the *E. coli* genes to obtain the parameters of the CRF model, i.e., the weights in the state pair dictionary and the state observation pair dictionary (Lafferty et al., 2001). The results of the IIS algorithm are shown in Fig. 2. All the delta weights of edges are less than 1e-6 (Fig. 2A) and all the delta weights of the vertices are less than 1e-6 (Fig. 2B), indicating that all parameters converge to the predefined range. Fig. 2C and 2D show the histograms for the weights of the edges and vertices, respectively. The values of most weights of the edges and vertices are around zero, however, the weights of a few edges and vbertices are far away from zero, such as the weight of TGCTAG is -19.26 and the weight of CTGLs is -3.17 (see the screen output of 3_get_delta_weights_of_dict.py for detail). To intuitively illustrate the effect achieved by parameter learning of the model, Fig. 2E shows the difference between the expected count and the empirical count as a percentage of the empirical count for each edge and Fig. 2F shows the difference between the expected count and the empirical count as a percentage of empirical count for each vertex. It can be seen that all of the errors are less than 1%. These results demonstrate that the parameter learning of the CRF model achieves the intended goal, which lays a solid foundation for the subsequent analysis.

**Fig. 2.**
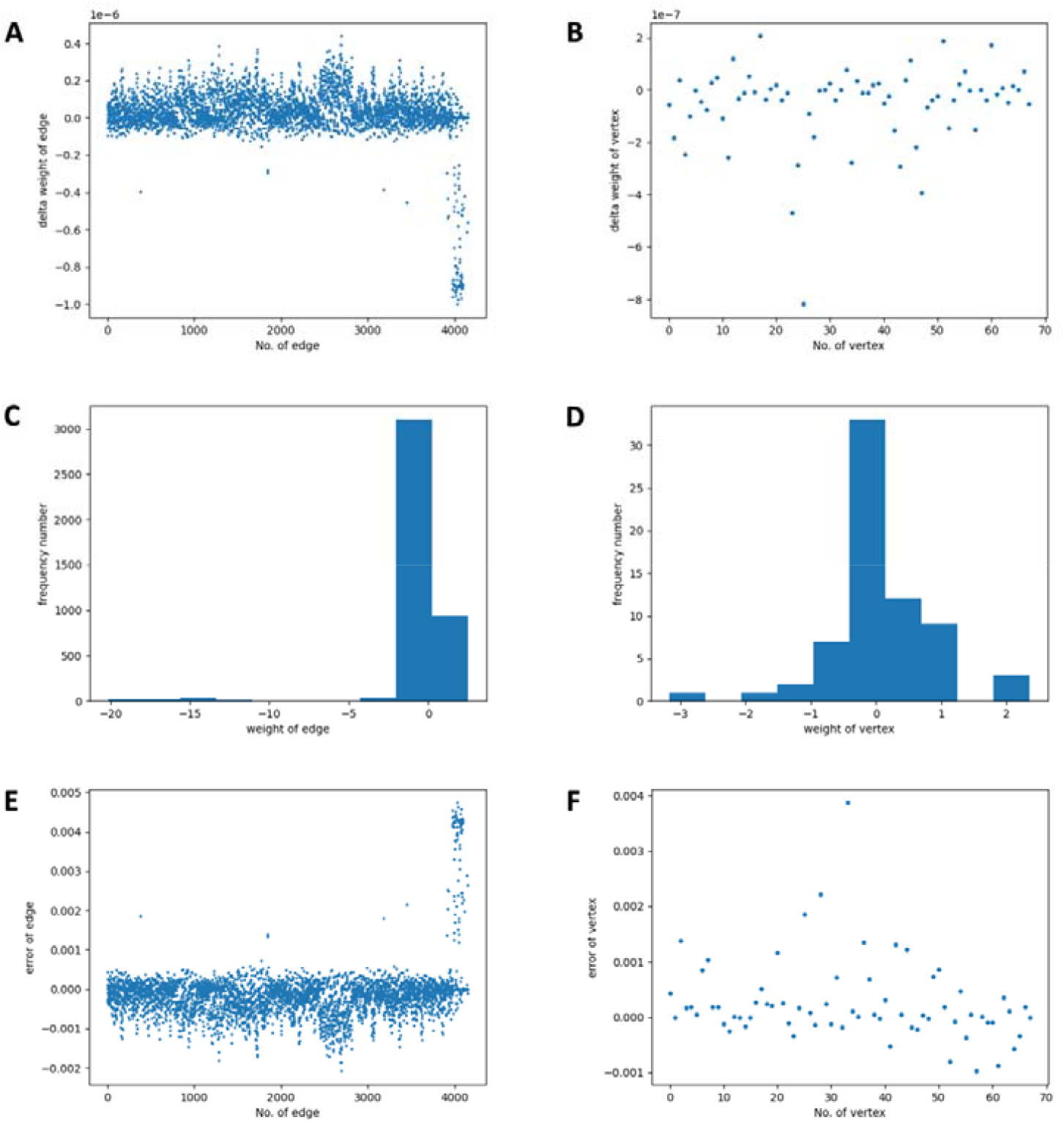
The learned parameters of conditional random field model. (A) The delta weights of the edges. (B) The delta weights of the vertices. (C) The weights of the edges. (D) The weights of the vertices. (E) The difference between the expected count and the empirical count as a percentage of the empirical count for each edge. (F) The difference between the expected count and the empirical count as a percentage of the empirical count for each vertex.

### Analysis of *E. coli* genes with CRF

We start by defining an indicator called the codon usage index for quantifying the codon usage of genes. In theory, many possible DNA sequences can encode a protein sequence of interest. For any two of these DNA sequences, such as DNA sequence 1 and DNA sequence 2, the first codons of the DNA sequence 1 and DNA sequence 2 may be the same, or different, and so on for the rest of the codons. If we count the number of situations where the two codons are the same and then divide by the total number of codons in DNA sequence 1, then we get the codon usage index for DNA sequence 1 and DNA sequence 2. For a given protein sequence, the mathematical expectation of the codon usage index for the two random DNA sequences can be calculated with the frequency of codon usage, or calculated with the frequency of codons as equally possible. Since the parameters of the CRF model have been obtained, the DNA sequence of max probability according to the Viterbi algorithm then is compared with the real gene sequence to get the codon usage index.

With the learned parameters of the CRF model, the Viterbi algorithm is used to analyze the codon usage indexes of the 4292 genes of *E. coli* (Fig. 3) (Viterbi, 1967). For most of the genes, the codon usage index between the real gene sequence and the max probability gene sequence is greater than the mathematical expectation of the codon usage index considering the frequency of codon usage and even greater than the mathematical expectation of the codon usage index with the frequency of codons as equally possible (Fig. 3A). The difference in the codon usage index is greater than 0 for nearly 4, 000 genes. The groups with the largest frequency number are within the range of 0.1 to 0.2, accounting for over 2, 000 genes (Fig. 3B). These results demonstrate that the real gene sequence tends to use codons in a way similar to how the DNA sequence of the max probability of the CRF model uses codons.

**Fig. 3.**
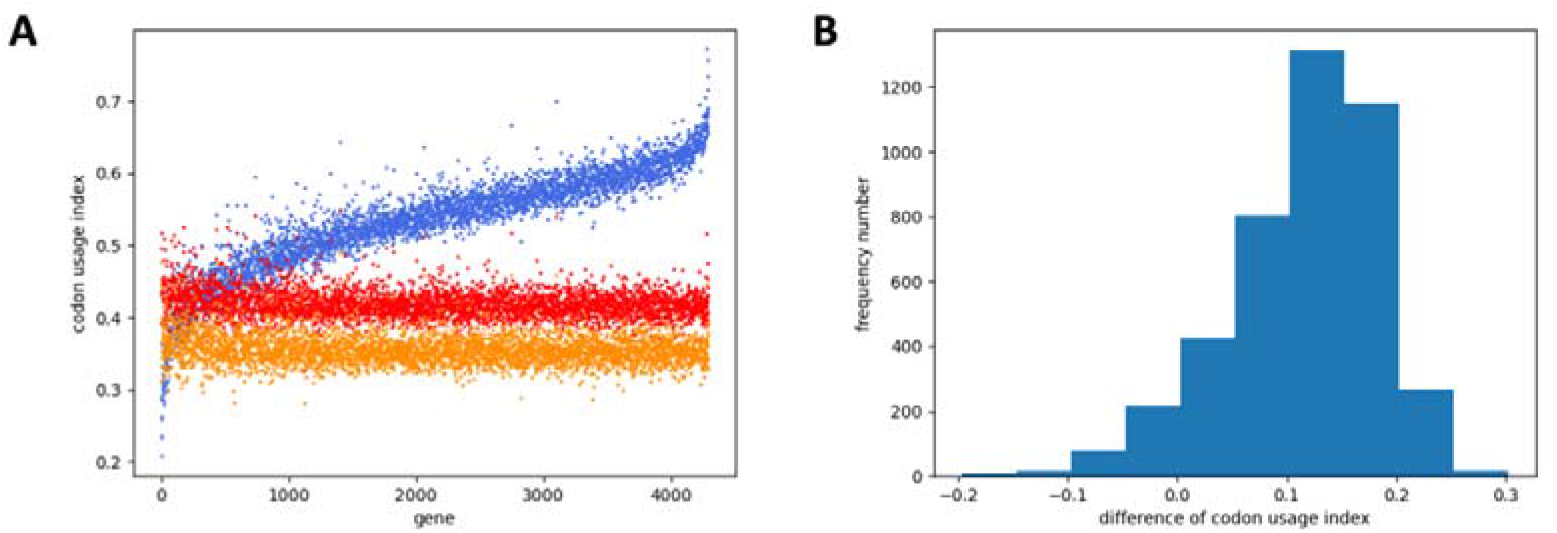
Codon usage analysis of *E. coli* genes with the conditional random field model. (A) The sorted codon usage indexes for *E. coli* genes. The orange dot shows the mathematical expectation of the codon usage index for two random DNA sequences corresponding to the given *E. coli* protein sequence with the frequency of codons as equally possible. The red dot shows the mathematical expectation of the codon usage index for two random DNA sequences corresponding to the given *E. coli* protein sequence, taking the codon usage frequency into consideration. The blue dot shows the codon usage index for the max probability DNA sequence and the real DNA sequence for the given *E. coli* protein sequence. The genes are sorted increasingly according to the value of the blue dot minus that of the red dot. (B) The frequency number of the differences between the values of the blue dot and the corresponding red dot in (A).

### Distribution of the empirical counts for the edges and vertices of *E. coli* genes

To investigate the distribution of the empirical counts for the edges and the vertices of *E. coli*, a random number of random genes are selected from *E. coli* genome to get the edge count and the vertex count. The Pearson product-moment correlation coefficient (PPMCC) is calculated for the edge counts of each sampling (containing a pair of samples) from the genes of *E. coli* genome (Fig. 4). For *E. coli*, the PPMCCs for edges are close to 1 and those for the vertices are even closer to 1. This data is unexpected and very interesting. Therefore, the genome sequences of three other bacteria are investigated and the results are similar to that of *E. coli*. These results demonstrated that for a random subset of the genes of some bacterial genomes, the distribution of edges follows a certain pattern, and the distribution of vertices follows a certain pattern as well. Both patterns are linear proportional, as the PPMCCs are very close to 1. This suggests that the codon usage of the genes of a bacterium might reach a stylistic consistency during a long time of evolution, and it is possible to learn approximately the parameters of the CRF model even if only a subset of the bacterium genes is available.

**Fig. 4.**
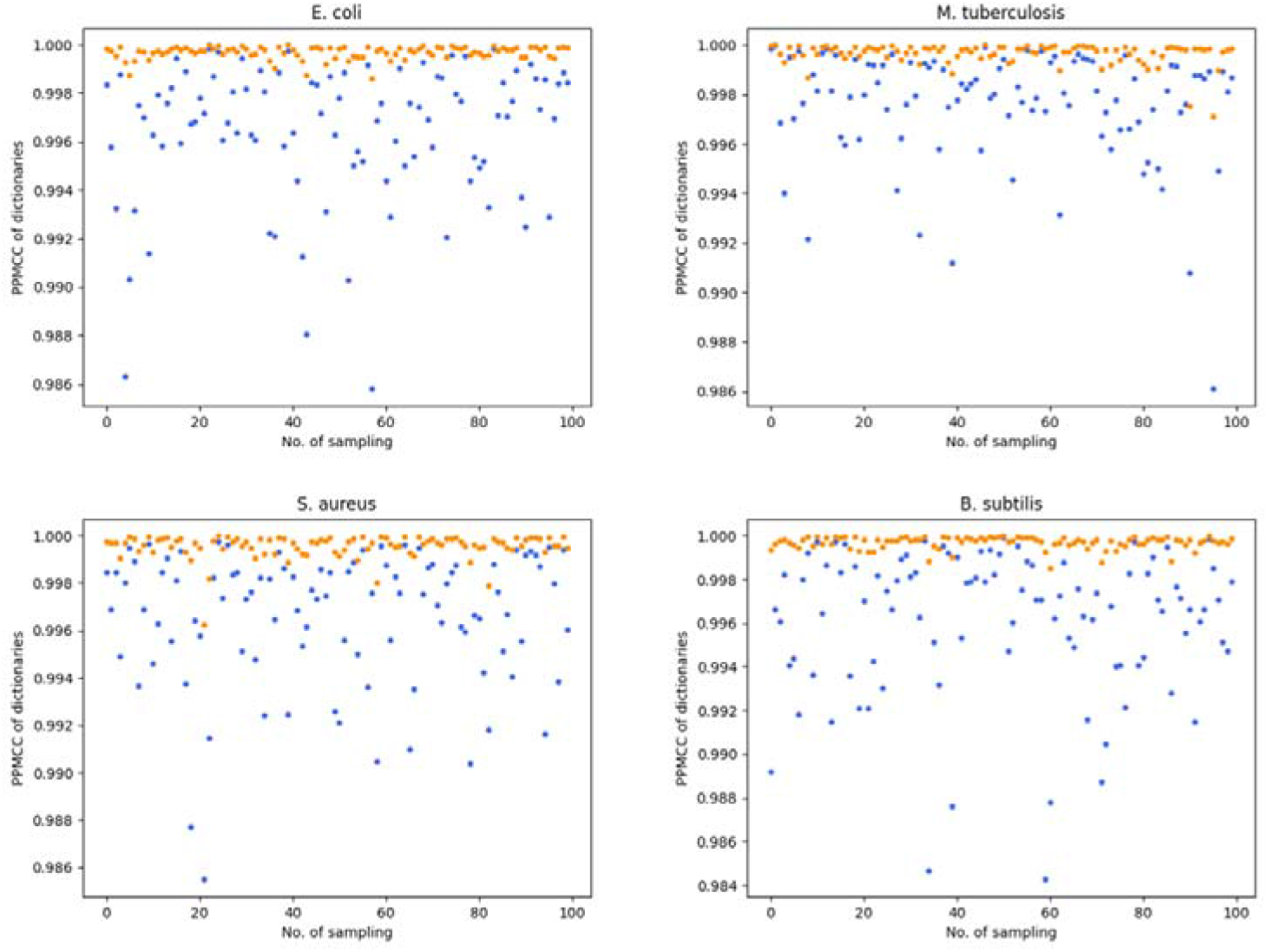
The distribution of edge and vertex of the genes for four bacteria. The name of the bacterium is shown on the title of each subplot. The blue dot shows the PPMCC for the edges of each sampling and the orange dot shows the PPMCC for the vertices of each sampling.

### Analysis of phage genes with CRF

The phages depend on host *E. coli* to complete their life cycles (Salmond and Fineran, 2015). Though they have no protein manufacturing machinery, the phages might have learned to acquire the rules of codon usage from their host during the process of evolutionary adaptation. To test this, the genomes of eight phages, including T1-T7 and lambda, are used for CRF analysis (Fig. 5). Except for the T7, the pattern of the codon usage indexes for the genes is similar to that of *E. coli* genes. More specifically, for most genes of the T1-T6 and lambda, the codon usage of the real gene sequence is similar to that of the max probability DNA sequence decoded by the CRF model of *E. coli*. However, the T7 genes have codon usage indexes close to the mathematical expectation, which means that the T7 has not learned enough from the host about the codon usage style. Possibly the reason is that the T7 is a virulent phage, which has not reached a balance with the host. These results demonstrate that phages can adapt to the stylistic codon usage of the host and also provide a new viewpoint for examining the relationship between the phages and *E. coli*.

**Fig. 5.**
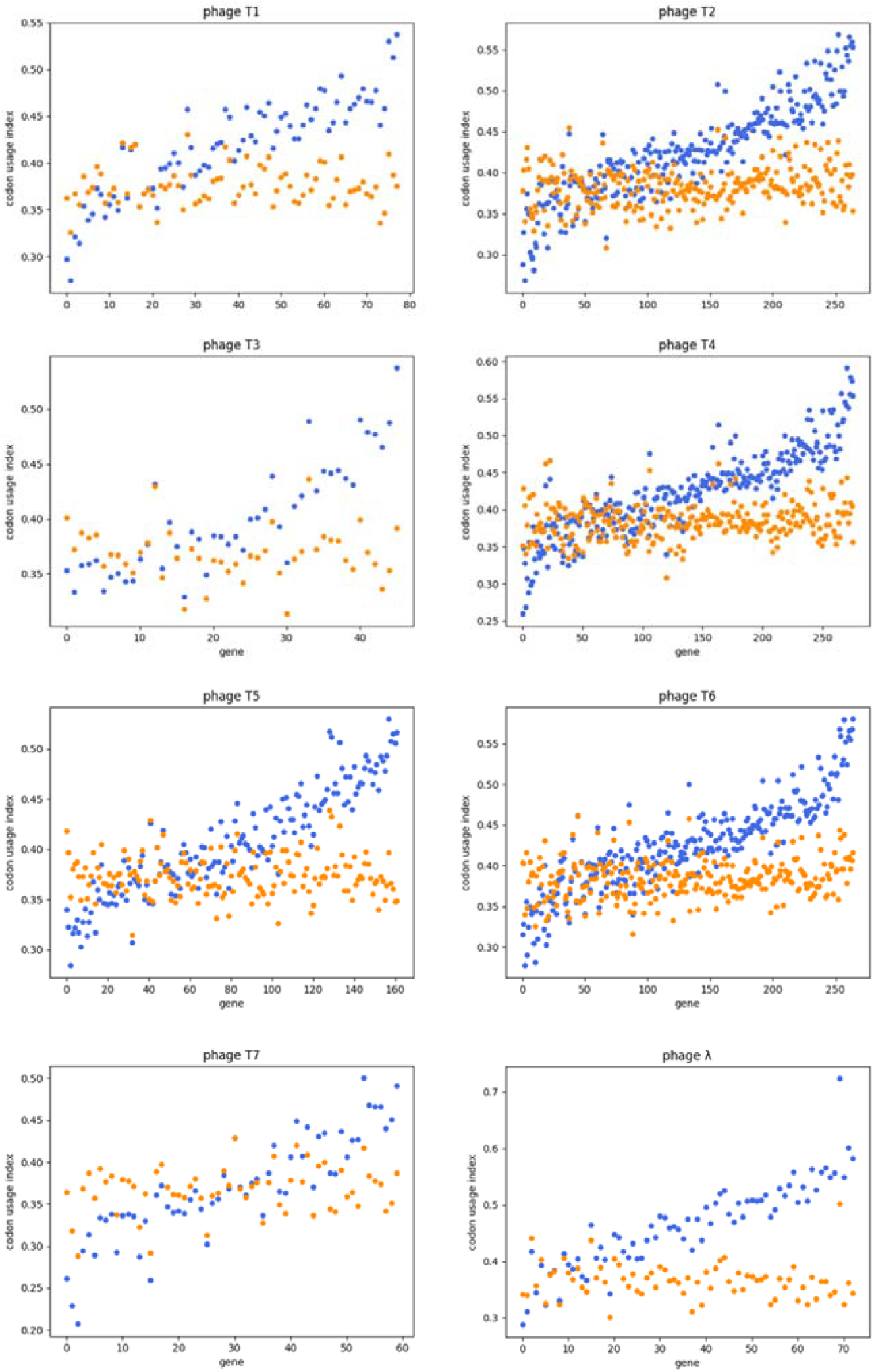
Codon usage analysis of phage genes with the conditional random field model. The name of the phage is shown on the title of each subplot. The orange dot shows the mathematical expectation of the codon usage index for one given gene sequence with the frequency of codons as equally possible. The blue dot shows the codon usage index between the max probability DNA sequence and the real DNA sequence. The genes are sorted in an increasing manner according to the value of the blue dot minus that of the orange dot.

### Analyzing codon optimization of the genes with CRF

To investigate whether the CRF model can be used in the codon optimization of the heterologous genes expressed in *E. coli*, three genes, i.e., IL-2 (Williams et al., 1988), TTFC (Makoff et al., 1989), and NF-1-334 (Hale and Thompson, 1998) are used for analysis. It has been reported that the soluble expression of IL-2, TTFC, and NF-1-334 are improved 16, 4, and 3 folds respectively via codon optimization (Gustafsson et al., 2004). The original sequences and the optimized sequences are prepared and analyzed with the *E. coli* CRF model (Fig. 6). For the original gene sequences, the codon usage indexes are near the mathematical expectation, suggesting that the codon usage of the original gene sequences is random (Fig. 6A). For the optimized gene sequences, the codon usage indexes are higher than the mathematical expectation, which means that the codon usage of the optimized gene sequences conforms to that of the *E. coli* genes (Fig. 6B). These results suggest that the CRF model can play a useful role in the codon optimization of the heterologous gene expression.

**Fig. 6.**
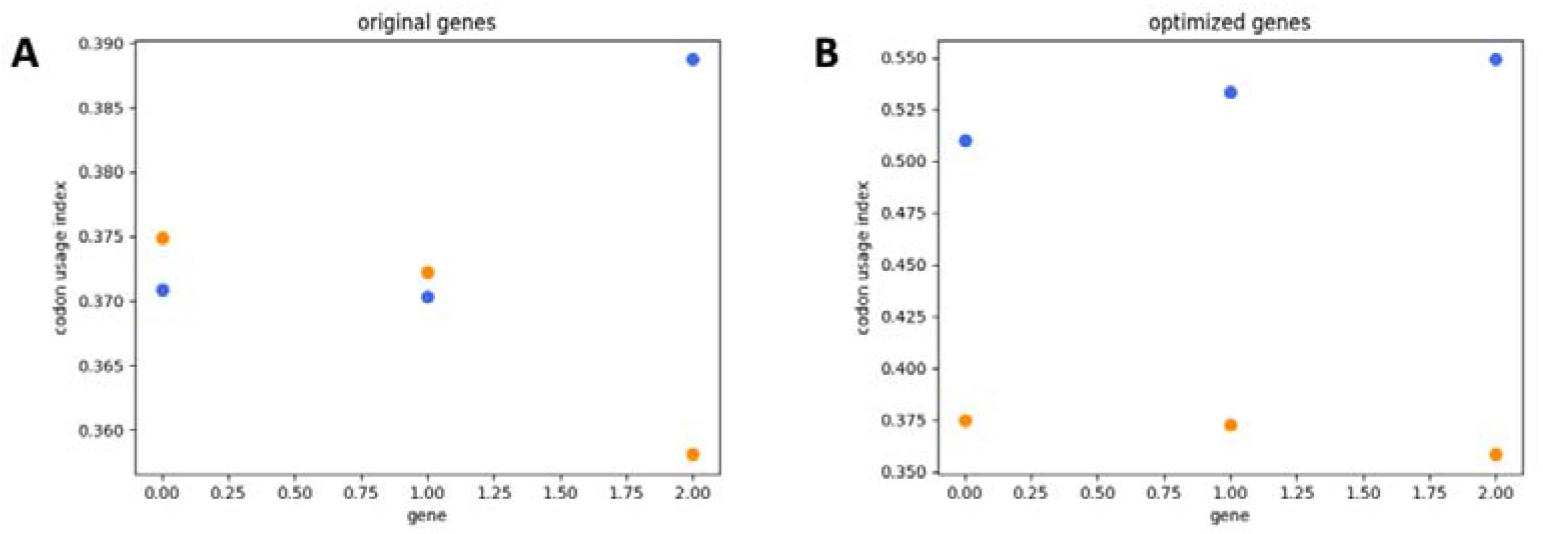
Codon usage analysis of three example genes with the conditional random field model. (A) Codon usage analysis of the original genes. (B) Codon usage analysis of the optimized genes. The orange dot shows the mathematical expectation of the codon usage index for one given gene sequence with the frequency of codons as equally possible. The blue dot shows the codon usage index between the max probability DNA sequence and the real DNA sequence. The genes are sorted increasingly according to the value of the blue dot minus that of the orange dot.

## Discussion

In this study, the codon usage of the protein-encoding genes is addressed with the CRF model to ensure that the model is structurally elegant and mechanistically interpretable. The results of the parameter learning can be explained clearly. Locally, a greater weight of the edge or the vertex indicates a greater possibility of the edge or the vertex. Globally, the possibility of a certain DNA sequence for the protein sequence in question is determined by all of its edges and vertices and their corresponding weights, in the context of all possible DNA sequences.

During the analysis of *E. coli* genes with the CRF model, it is shown that the codon usage of the real gene sequence does not conform 100% to the DNA sequence of max probability. This may be explained plainly. The CRF model is like the voting system of our society. Each gene votes for the parameters of the CRF model with its edges and vertices. The voting system counts the number of each type of edge and each type of vertex and subsequently releases a set of weight-assigned parameters that both stands for the collective choice internally (genes are interacting via voting) and reflects the actual situation externally. Most genes may be happy to find it consistent with their ballots more or less, while the rest may notice that the result does not match well with their choices, or even runs counter occasionally. However, the weights of the parameters are determined jointly by all genes, to satisfy all genes instead of some specific genes. Thus, the max probability DNA sequence calculated with such a weight system fits largely rather than completely to the real DNA sequence.

In sum, the application of the CRF model to the *E. coli* codon usage helps us understand the codon usage systematically and quantitatively. It is useful for the codon optimization of the heterologous gene expression and may potentially inspire the transgene, mRNA vaccine design, and synthetic biology.

## Materials and methods

### Computer hardware and software

The running environment includes MacBook Air (M1, 2020), macOS Monterey 12.3.1, and Python 3.10.4 (Van Rossum and Drake, 1995) with packages numpy 1.23.1 (Harris et al., 2020), mpmath 1.2.1 (Johansson et al., 2021), pandas 1.4.3 (McKinney, 2010), biopython 1.79 (Cock et al., 2009), and matplotlib 3.5.2 (Hunter, 2007). The improved iterative scaling (IIS) program 7_improved_iterative_scaling.py needs hardware double Intel Xeon E5-2696 v3 CPUs with each CPU 18 cores/36 threads and software CentOS Linux release 7.9.2009 (Core) with conda 4.13.0 (Anonymous, 2020). The Python 3.10 environment is created in conda and the required package mpmath 1.2.1 is pip installed. The IIS program 7_improved_iterative_scaling.py (may need to check and modify the value of the variable sub_process_count in line 324 to a proper value such as 8 or 72 or other depending on the CPU before running) runs over 50, 000 cycles (each cycle taking 38 seconds) before converging to max delta weight less than 1e-6 for all parameters of the CRF. All other programs can be run with MacBook Air quickly, in seconds or a few minutes.

### Preparation of gene data for *E. coli*

The codes and data for this section are in Supplementary Material S1. The genome fasta file (MG1655.fasta) of *Escherichia coli* str. K-12 substr. MG1655 is downloaded from https://www.ncbi.nlm.nih.gov/nuccore/U00096.3?report=fasta. The gene information of the proteins (proteins_167_161521.csv) of *Escherichia coli* str. K-12 substr. MG1655 is downloaded from https://www.ncbi.nlm.nih.gov/genome/browse/#!/proteins/167/161521%7CEscherichia%20coli%20str.%20K-12%20substr.%20MG1655/chromosome/. The script 1_find_gene_id_duplications.py is used to look for the genes with the same IDs (944843, 945105, 946106, 945687, 946401, 947077, 947684, 63925658, 949122) in the proteins_167_161521.csv, which are modified to id-2 or id-3 and treated as new genes. The 2_get_genes.py is used to extract the gene sequences from the genome sequence and stored in the folder named gene_files. The 3_check_genes.py is used to check whether each gene has a complete coding frame, i.e., having a start codon and a stop codon, and to exclude those genes of which the base numbers are not the folds of 3 (945105-2, 946106-2, 947369, deleted from gene_files). The 4_check_proteins.py is used to check whether there is any stop codon in the middle of the gene sequence (excluding the first and the last codons). Those genes with at least one stop codon in the middle of the sequence (946035, 948394, 948584) are deleted from gene_files. After all these *E. coli* gene data preparation and data cleaning, a total of 4292 genes are obtained.

### Presentation of the CRF model

The structure of the probabilistic graph model CRF is defined with the keys of the Python dictionaries. Reference (Alberts, 2015) is used to prepare the file 0_codon_table_raw.txt, which records the codon dictionary. There are two dictionaries for describing CRF model in this study, including the state pair dictionary and the state observation dictionary. The state pair dictionary contains the weight, the empirical count, and the expected count for each edge of the CRF model, and the state observation dictionary contains the weight, the empirical count, and the expected count for each vertex of the CRF model. The 1_build_dictionaries.py is used to obtain these two dictionaries. Along with three other dictionaries produced by the 1_build_dictionaries.py, they are all used for parameter learning of the model. The codes and data for this section are in Supplementary Material S2.

### Parameter learning and analysis of CRF

The 1_calculate_empirical_features.py is used to get the counts of the state pairs and state observation pairs in the genes of *E. coli* genome, which are stored in the state pair dictionary and the state observation dictionary, respectively. The 2_improved_iterative_scaling.py is used for the iterative optimization of the weights of the state pair dictionary and the state observation dictionary until the max_delta_weight < 1e-6. The folder zzz_CRF_weights_final stores the two dictionaries with learned weights. The codes and data for this section are in Supplementary Material S3.

Then the two dictionaries are analyzed with the programs in Supplementary Material S4. The 1_compare_two_dicts.py is used to compare the empirical counts before and after IIS optimization. The 2_get_numbers_of_edges_and_vertices.py is used to get the counts of the edges (4156 types) and vertices (70 types). The 3_get_delta_weights_of_dict.py is used to get the delta weights of the state pair dictionary and the state observation dictionary. The 4_get_errors_between_expected_and_empirical_for_dicts.py is used to get the errors between the empirical and expected counts of the state pair dictionary and the state observation dictionary.

### Analysis of *E. coli* genes with CRF

The codes and data for this section are in Supplementary Material S5. The 1_get_proteins_from_genes.py is used to translate the *E. coli* genes to proteins, which are stored in the folder named protein_files. The 2_get_gene_max.py is used to get the DNA sequence of max probability according to the Viterbi algorithm for each protein sequence, which is stored in the folder named gene_max_files. The 3_get_codon_frequency3.py is used to get the frequency of codon usage in *E. coli*. The 4_get_codon_identity_for_each_gene.py is used to get the codon usage indexes for each protein, including the codon usage index between the real DNA sequence and the DNA sequence of max probability, the mathematical expectation of the codon usage index for each protein with the frequency of codon usage, and the mathematical expectation of the codon usage index for each protein with the frequency of codons as equally possible. The results are sorted with the codon usage index between the real DNA sequence and the DNA sequence of max probability minus the mathematical expectation of the codon usage index for each protein with the frequency of codon usage, and stored in the file ratios_of_identical_codons_of_genes.txt.

### Analyzing the distribution of the empirical counts for the edges and vertices of *E. coli* genes

The codes and data for this section are in Supplementary Material S6. The 1_build_dictionaries.py is used to produce the state pair dictionary and the state observation dictionary, both with zero count for each empirical value of the edges or vertex. The 2_analyze_empirical_features.py is used to select a random number of random genes from *E. coli* genes and count the edges and vertices and the results are stored in dictionaries. For two times of such random selection and counting, the Pearson product-moment correlation coefficient (PPMCC) can be calculated for each pair of edge dictionaries and for each pair of vertex dictionaries. Three other bacteria are investigated as well. The mutifasta file of the *Mycobacterium tuberculosis* H37Rv genome is downloaded from https://www.ncbi.nlm.nih.gov/nuccore/NC_000962.3. The mutifasta file of the *Staphylococcus aureus* subsp. aureus NCTC 8325 is downloaded from https://www.ncbi.nlm.nih.gov/nuccore/NC_007795.1. The mutifasta file of the *Bacillus subtilis* subsp. subtilis str. 168 is downloaded from https://www.ncbi.nlm.nih.gov/nuccore/NC_000964.3. The 2_get_single_genes_from_multifasta.py is used to get the single genes from a multifasta file and stored in the folder named gene_files. The 3_check_genes.py is used to filter those genes that are ineligible for analysis (delete if found). The 4_analyze_empirical_features.py is used to get the PPMCC of the edges and the PPMCC of the vertices.

### Analysis of phage genes with CRF

The codes and data for this section are in Supplementary Material S7. The phages co-evolute with the host *E. col*i, which have no translation machinery to produce their proteins and must depend on host *E. coli* to accomplish the task of manufacturing proteins. The genes of the phages T1-T7 and lambda are analyzed with the *E. coli* CRF model to see if they conform to the codon usage style of *E. coli*.

The NCBI accession numbers of the Enterobacteria phages are T1: NC_005833.1, T2: AP018813.1, T3: NC_047864.1, T4: NC_000866.4, T5: NC_005859.1, T6: NC_054907.1, T7: NC_001604.1, lambda: NC_001416.1. The multifasta file of each genome is downloaded with the corresponding accession number from NCBI. The 1_get_single_genes_from_multifasta.py is used to get each gene sequence from the phage genome. The 2_get_proteins_from_genes.py is used to get the protein sequence for each gene. The 3_get_gene_max.py is used to get the gene sequence of max probability for each protein sequence. The 4_get_codon_identity_for_each_gene.py is used to get the codon usage indexes for each protein, including the codon usage index between the real DNA sequence and the DNA sequence of max probability, and the mathematical expectation of the codon usage index for each protein with the frequency of codons as equally possible. The results are sorted with the codon usage index between the real DNA sequence and the DNA sequence of max probability minus the mathematical expectation of the codon usage index for each protein with frequency of codons as equally possible, and stored in the file ratios_of_identical_codons_of_genes.txt. For phage T4, two genes containing characters other than ATCG and the two gene files are deleted before further analysis.

### Analyzing codon optimization of the genes with CRF

The genes IL-2 (Williams et al., 1988), TTFC (Makoff et al., 1989), and NF-1-334 (Hale and Thompson, 1998) from reference (Gustafsson et al., 2004) are analyzed with CRF. The above references provide the original DNA sequences and the optimized gene sequences, which are convenient for comparison with CRF. The codes and data for this section are in Supplementary Material S8. To make the gene sequence suitable for analysis, the ATG is added to the 5’ end for the IL-2 sequence, and the ATG is added to the 5’ end and the TAA is added to the 3’ end for the TTFC sequence. The 1_get_proteins_from_genes.py is used to get the protein sequence for each gene sequence. The 2_get_gene_max.py is used to get the DNA sequence of max probability for each protein sequence. The 3_get_codon_identity_for_each_gene.py is used to get the codon usage indexes for each protein, including the codon usage index between the real DNA sequence and the DNA sequence of max probability, and the mathematical expectation of the codon usage index for each protein with the frequency of codons as equally possible. The results are sorted with the codon usage index between the real DNA sequence and the DNA sequence of max probability minus the mathematical expectation of the codon usage index for each protein with the frequency of codons as equally possible, and stored in the file ratios_of_identical_codons_of_genes.txt.

## Acknowledgments

We thank Mr. Sheng Lin for his help with the hardware and software of the Linux CentOS server.

## Additional information

### Data availability

All Python codes for analyzing the data, i.e., Supplementary Material S1 to S8 are available at https://github.com/shiqiang-lin/crf_e_coli.

